# PhenoScreen: A Dual-Space Contrastive Learning Framework-based Phenotypic Screening Method by Linking Chemical Perturbations to Cellular Morphology

**DOI:** 10.1101/2024.10.23.619752

**Authors:** Shihang Wang, Qilei Han, Weichen Qin, Lin Wang, Junhong Yuan, Yiqun Zhao, Pengxuan Ren, Yunze Zhang, Yilin Tang, Ruifeng Li, Zongquan Li, Wenchao Zhang, Shenghua Gao, Fang Bai

## Abstract

Phenotypic drug discovery (PDD) screens compounds in cellular models that represent disease-relevant phenotypes, offering a compelling alternative to traditional target-based approaches. Unlike conventional methods, where compounds act on a single predefined target, PDD identifies compounds capable of exerting therapeutic effects through multiple targets and mechanisms. This makes PDD particularly valuable for discovering first-in-class drugs, especially for diseases with poorly understood molecular mechanisms or those lacking validated therapeutic targets. By enabling broader exploration of biological systems and uncovering multi-target drugs (polypharmacology), PDD provides a powerful strategy for tackling complex diseases. In this study, we introduce PhenoScreen, a deep learning framework designed to advance PDD by utilizing large-scale compound-phenotype association data. Through contrastive learning, PhenoScreen connects chemical space with cellular morphological profiles, allowing for accurate prediction of compound-induced phenotypic changes. PhenoScreen can also accurately identfiy lead compounds which could induce user defined phenotypic shift but more novel scaffolds using different levels of phenotypic information reflected by diverse compounds. The model was validated across multiple screening tasks and successfully predicted active compounds inducing user-specified phenotypes with varying inhibitory effects in the osteosarcoma phenotypic model. Further, other than showing effectiveness to osteosarcoma, our experiments also showed that PhenoScreen demonstrated strong generalization to rhabdomyosarcoma, and the active compound we screened had an IC_50_ of up to 1.842 μM, suggesting its ability to capture key phenotypic features shared across cancer cells. These results underscore PhenoScreen’s potential to accelerate drug discovery by identifying novel therapeutic pathways and increasing the diversity of viable drug candidates. PhenoScreen is accessible online via our group’s web server for compound virtual screening at https://bailab.siais.shanghaitech.edu.cn/services/PhenoScreen/, and the source codes are available at https://github.com/Shihang-Wang-58/PhenoScreen.

## Introduction

Phenotypic drug discovery (PDD) is a classic and powerful strategy in drug development.^1–3^ Traditionally, drug discovery methods relied heavily on target-based drug discovery (TDD), whose focus was on identifying compounds that interact with a specific biological target.^4^ However, phenotype-based approaches do not require prior knowledge about the target. Instead, PDD focuses on identifying compounds that elicit desired phenotypic effects in biological systems, such as cell morphology changes, disease state reversal, or specific signaling pathway modulation.^5^ This allows the identification of potential therapeutic compounds that act on novel or multiple targets, making PDD particularly valuable for diseases with unknown molecular mechanisms or limited treatment options. A recent retrospective analysis highlighted that most first-in-class drugs originated from PDD rather than TDD, highlighting the importance of PDD in discovering innovative therapeutics for complex diseases.^4,6^ PDD offers several advantages over TDD.^4^ First, it provides a broader exploration of biological systems and can identify multi-target drugs (polypharmacology), which often enhance therapeutic efficacy. Furthermore, PDD is well-suited for discovering first-in-class drugs by targeting unknown or novel mechanisms, which is crucial for addressing complex or poorly understood diseases. However, PDD also faces many challenges, including the difficulty in target identification and the high cost of large-scale screening efforts. Advances in high- content imaging, such as the Cell Painting technique, have provided new tools for capturing cellular phenotypes, but real-world screening is still resource-intensive, and efficient computational methods are needed to process and analyze phenotypic data on a large scale.^7^

AI, particularly deep learning, has been pivotal in analyzing complex phenotypic data, including high-content screening (HCS) images, transcriptomic profiles, and multi-omics datasets, enabling the discovery of potential drug candidates even without well-defined molecular targets. Employing diverse computational biology methods to aid early drug discovery can help researchers explore a wider chemical space that is currently beyond the reach of experimental methods.^8^

AI-driven approaches have the potential to accelerate drug discovery by integrating phenotypic information with molecular representation, enabling faster and more accurate predictions of compound efficacy.^9^ For example, Tong et. al proposed the TranSiGen model based on variational autoencoder to represent phenotypic information related to the cellular environment and compound effects, as well as phenotype-based drug redirection.^10^ Ni et. al proposed the PertKGE model based on the knowledge graph, and applied it to the phenotype- and target-based drug discovery process, which led to the discovery of a variety of novel hit compounds.^11^ Recent studies have also shown that cellular morphological profiles can be complementary to compound structure data in predicting drug efficacy. Integrating these phenotypic data with machine learning and deep learning techniques has become an emerging field of research in drug discovery.^1^ For instance, the Broad Institute’s Cell Painting technique, which captures high-content cellular images to analyze cell morphology, has proven effective in revealing drug mechanism of action (MOA), predicting bioactivity, and assessing toxicity. ^1,12–18^ The Klambauer lab initially used high-throughput microscopy imaging data and convolutional neural networks to predict the results of biological assays for different compounds.^19^ Haslum et.al have developed a deep learning based high-throughput cell image analysis method to predict bioactivities of different compounds.^15^ Recently, Dee et.al applied the SwinV2 model to predict the mechanism of action of 10 kinase inhibitor compounds directly from Cell Painting images.^18^ When integrated cellular morphological profiles with AI models, this approach can enhance molecular representation by incorporating cellular phenotypic responses alongside structural data, offering a more holistic view of a compound’s biological effects. In recent years, the Broad Institute has released multiple Cell Painting datasets, completed in collaboration with various pharmaceutical companies and academic institutions. These datasets contain a vast amount of cellular microscopy images and omics analyses, covering hundreds of thousands of compounds as well as perturbations from gene knockouts and overexpressions,^20,21^ providing an opportunity to explore powerful deep learning framework to develop phenotypic-based drug screening or even generation methods.

The chemical space is vast, making it impossible to fully explore in search of bioactive molecules for specific bioassays. This remains one of the most challenging issues that needs to be addressed. Some studies are starting to emerge to explore the strategies to combine molecular structure with other characteristics related to phenotypic bioactivities to improve virtual screening.^19^ For example, CLOOM and MIGA have employed contrastive learning to combine molecular structures and cellular phenotypes, demonstrating improved performance in predicting biological activity and mechanisms of action.^16,17^ Overall, the application of machine learning and deep learning techniques in morphological analysis has greatly enriched phenotypic drug discovery, ranging from identifying the MOA of small molecules, optimizing lead compounds and predicting toxicological effects. However, these methods often depend heavily on high-quality cell images, which are often scarce, and may struggle to generalize well across different cell types. According to our limited knowledge, no experimentally validated computational method for phenotypic screening has been reported to date.

In this study, we propose a multimodal molecular representation model, PhenoScreen, based on our specifically designed dual-space contrastive learning framework to bridge the gap between molecular structure and phenotypic information. By jointly training molecular and image encoders, PhenoScreen integrates both chemical structure and cellular perturbation data, enhancing its capability to predict compound efficacy across multiple targets and disease models. Through extensive testing, validation and application studies, PhenoScreen has been demonstrated to have superior performance in virtual screening tasks and has been validated in different tumor cell lines, suggesting its potential to accelerate drug discovery by combining molecular properties and phenotypic effects in a unified framework, and to generalize to different drug discovery scenarios.

## Results and Discussions

### Dual-space Contrastive Learning Framework Improves Molecular Representation Learning

In this study, we proposed PhenoScreen, which is a dedicatedly designed dual-space contrastive learning model, and integrates molecules chemical space with their perturbated cellular morphology space, as illustrated in Fig. 1A. This representation model consists of four main modules: two feature extraction modules, the Molecule Graph Encoder and the Cell ViT Encoder; a primary molecule-molecule contrastive learning module; and a step-wise molecule-image contrastive learning module. To evaluate the PhenoScreen model, we performed modular ablation on two recognized target based virtual screening benchmark test sets, DUD-E and LIT-PCBA,^22,23^ and evaluated the overall active molecule screening ability (*AUROC*) and early enrichment ability (EF1%) of PhenoScreen, which helps to understand the specific contributions of the strategies used in constructing the PhenoScreen model.

**Fig. 1.**
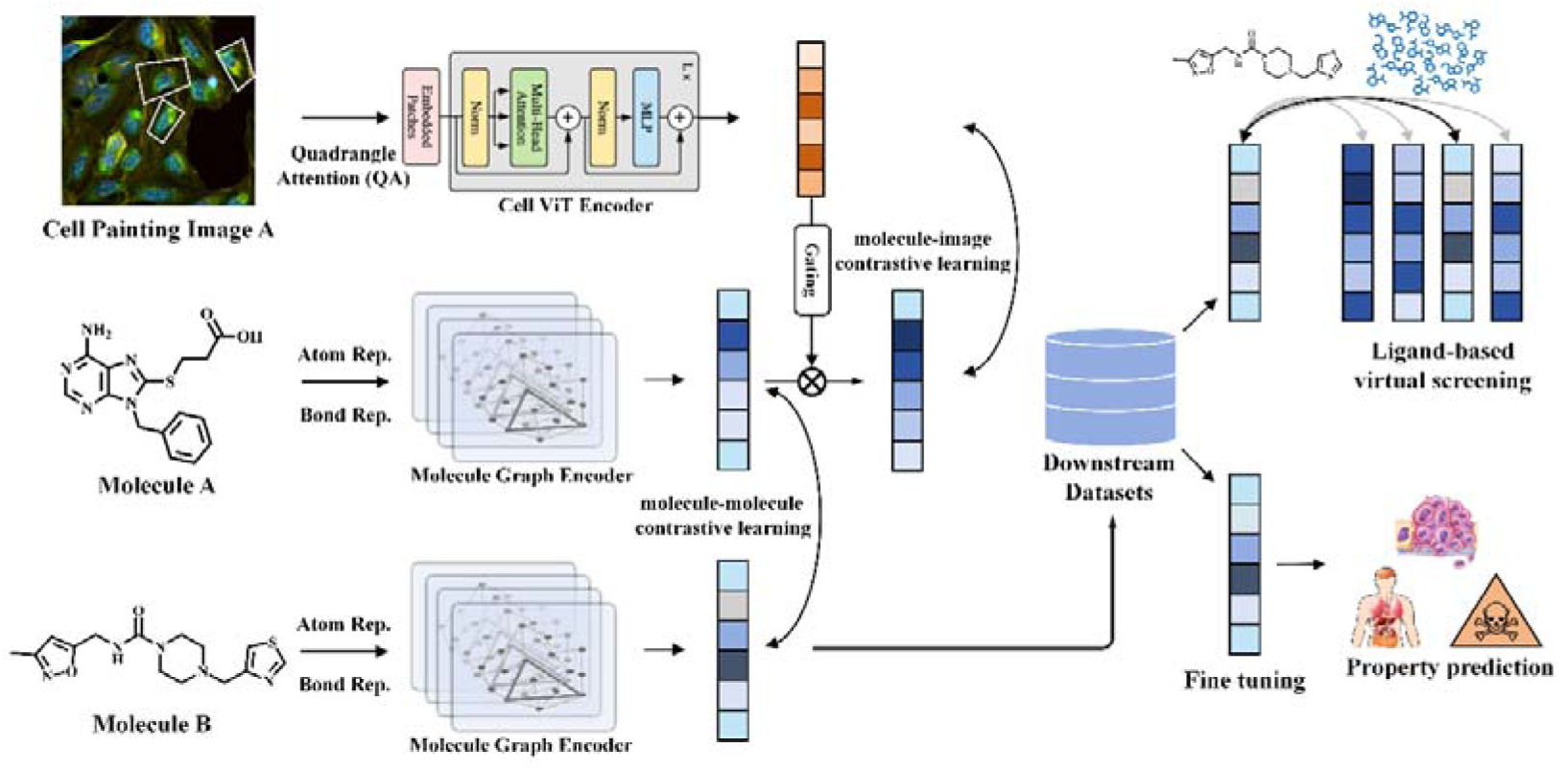
An illustration of the designed architecture for PhenoScreen. PhenoScreen consists of a molecule graph encoder, a Cell ViT Encoder, a primary molecule- molecule contrastive learning module; and a step-wise molecule-image contrastive learning module. This representation model can be then seamlessly integrated with a variety of drug design tasks, including zero-shot tasks such as virtual screening for active compounds, ADMET prediction, and other related applications.

We analyzed the contributions of molecule-image contrastive learning module, molecule-molecule contrastive learning module, and gating mechanism. As shown in Table 1, the version of PhenoScreen supplemented with cell image information (molecule-image contrastive learning module) outperformed models that lacked this information across all metrics, indicating that cell image information indeed enhances molecular representation. The screening results of active molecules for each target in LIT-PCBA are shown in Tables 2 and Tables 3. Notably, the performance of the model significantly declined when dual-space contrastive training was not utilized (without molecule-molecule contrastive learning module). This decline could be attributed to changes in the molecular encoder’s parameters during contrastive learning, as it aligns the molecule embeddings with image embeddings, potentially losing its original encoding ability. Thus, joint training maintains the molecular encoder’s intrinsic capability to effectively encode molecules while supplementing it with molecular activity information from cell images, resulting in improved representation performance of PhenoScreen. Additionally, removing the gating mechanism during contrastive training also led to a loss in performance to some extent. These results demonstrate the necessity of each module in PhenoScreen.

**Table 1.**
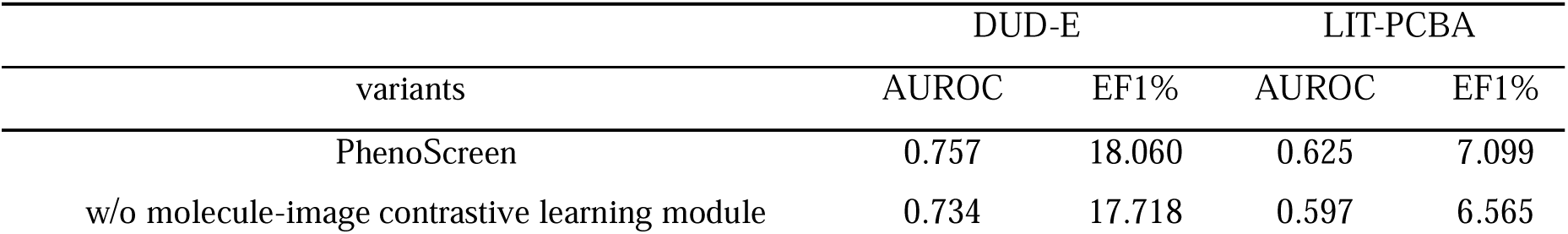
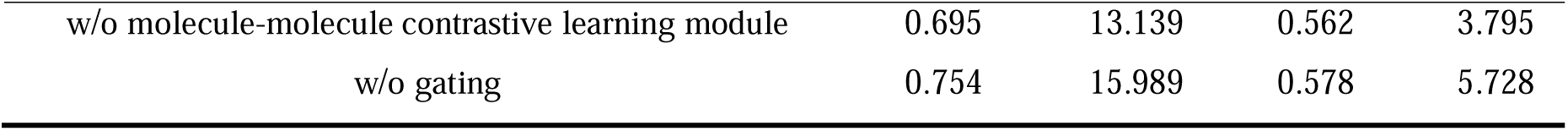
Performance of the PhenoScreen variants on DUD-E and LIT-PCBA.

| variants | DUD-E |  | LIT-PCBA |  |
| --- | --- | --- | --- | --- |
|  | AUROC | EF1% | AUROC | EF1% |
| PhenoScreen | 0.757 | 18.060 | 0.625 | 7.099 |
| w/o molecule-image contrastive learning module | 0.734 | 17.718 | 0.597 | 6.565 |
| w/o molecule-molecule contrastive learning module | 0.695 | 13.139 | 0.562 | 3.795 |
| w/o gating | 0.754 | 15.989 | 0.578 | 5.728 |

### Examining the Role of the Cell ViT Encoder in Enhancing Image Representation

Additionally, we evaluated the impact of different image encoders on the model’s contrastive training, as illustrated in Fig. 2. When using ResNet18 and ViT as image encoders, the different images corresponding to the same molecule were not effectively clustered together after encoding and dimension reduction. However, after incorporating Qformer Attention, multiple cell images corresponding to the same compound exhibited a much tighter distribution in the feature space. This indicates that the Cell ViT Encoder allows the image encoder to more accurately extract cellular features from cell images, enhancing the overall efficacy of the encoding process.

**Fig. 2.**
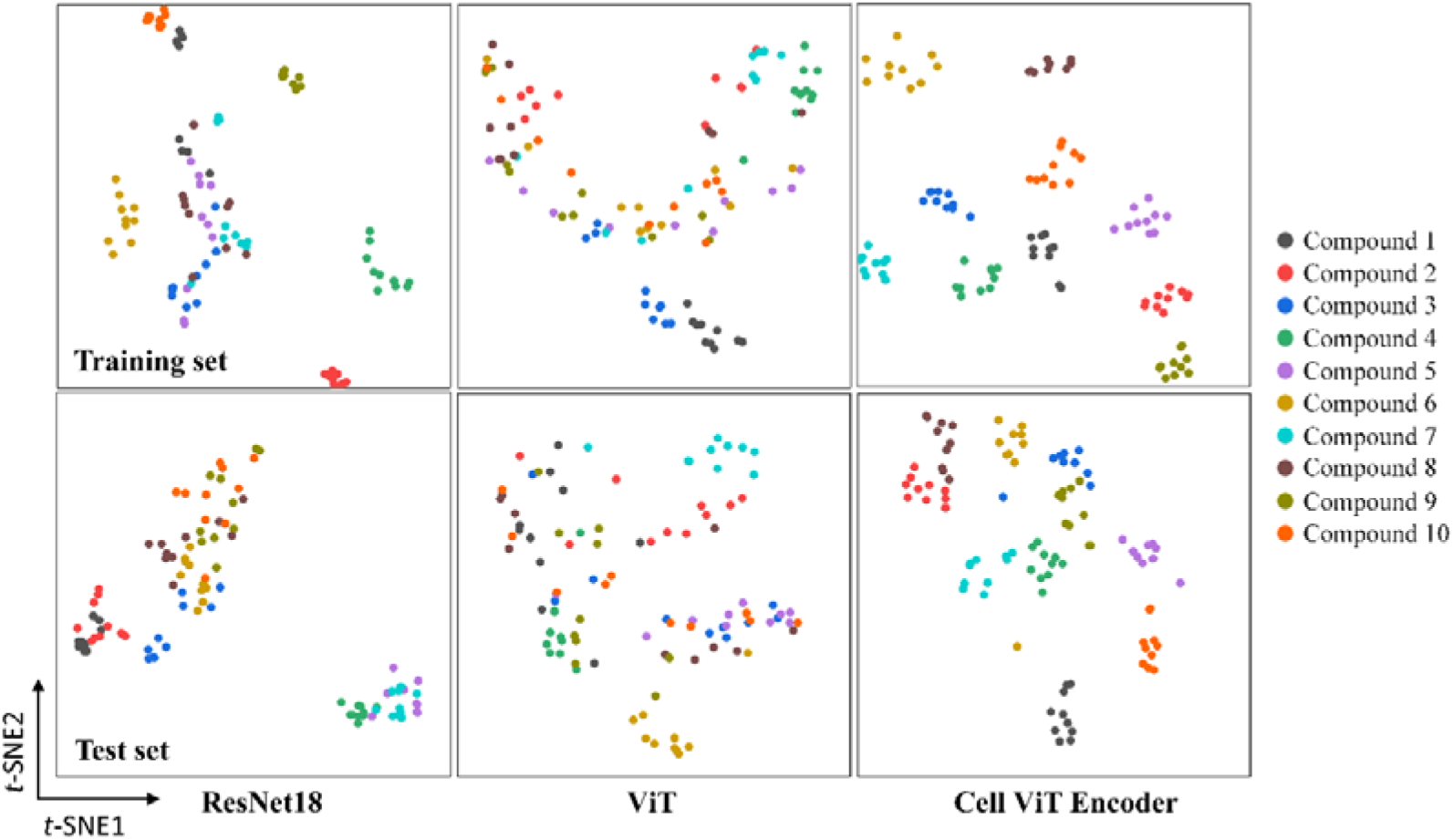
Visualization of Image Encoder Embedding Clusters. Cellular images of the same molecule are effectively clustered together with PhenoScreen encoding, demonstrating superior performance compared to other encoders such as ViT and ResNet18.

### PhenoScreen Surpasses Baseline Models in Phenotype-Based Virtual Screening

The phenotype information of cell images is included in the molecular image contrastive learning module and plays an important role in the PhenoScreen model. Considering that cellular phenotypes generally reflect phenotypic effects of molecules in biological systems or at the level of multi-target modulation. Therefore, we attempted to explore how cellular phenotype information complements the molecular representation capabilities of PhenoScreen. To this end, we collected 73 cell phenotype activity experimental datasets from the National Cancer Institute Development Therapeutics Program (NCI/DTP). These datasets come from 73 different cell lines of 10 types of cancer, including breast cell line, central nervous system cell line, colon cell line, leukemia cell line, melanoma cell line, non-small cell lung cell line, ovarian cell line, prostate cell line, renal cell line, and small cell lung cell line. The average number of test molecules per dataset is 43K, and the average number of active molecules is 2.8K. Given that cell activity may be influenced not just by a single target but also by the synergistic effects of multiple targets, this task presents greater challenges.

We selected eight different molecular fingerprints, an our group previously developed conformational ensemble-based molecular representation method GeminiMol, and PhenoScreen for benchmark testing comparison. During the testing process, all molecules were independently encoded by the corresponding method, and the activity of molecules against various targets was distinguished by calculating the similarity between query molecules and reference molecules. Considering that changes in cell phenotype may be influenced by multiple mechanisms of action, we randomly select multiple reference molecules for each biological assay. Finally, we calculated the average *AUROC* of these methods across all cell lines included in each type of cancer. As illustrated in Fig. 3 and Table S1, PhenoScreen showed outstanding performance in screening active molecules across all ten types of cancer cell lines, especially compared to the GeminiMol model which only contains molecule structure information for training, indicating the enhancement of the molecular encoder by cell phenotype information.

**Fig. 3.**
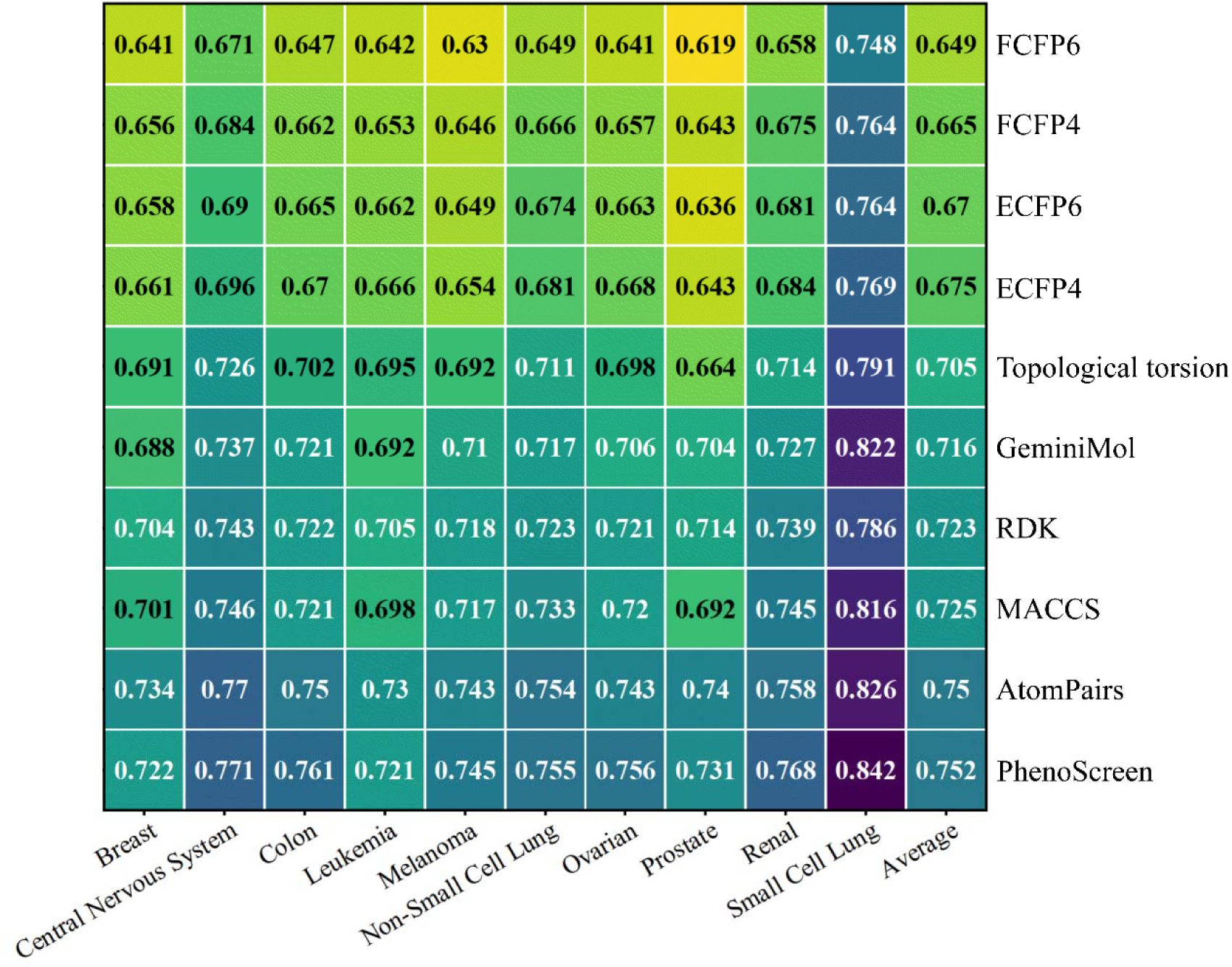
Performance of different methods in screening active molecules across 73 different cell lines of 10 types of cancer. The horizontal axis represents ten different types of cancer. The vertical axis represents the different types of molecule representation methods. The values in each square of the heatmap represent the average *AUROC* value of active molecule screening for this method in all cell lines of the denoted type of cancer.

### PhenoScreen Also Excels in Target-Based Screening of Active Compounds

Meanwhile, we tested the performance of the method over the two important virtual screening tasks in drug design: DUD-E and LIT-PCBA.^22,23^ DUD-E contained 102 targets and LIT-PCBA contained 15 targets for evaluating the performance of the virtual screening algorithms. All molecules were independently encoded by the PhenoScreen model, and the activity of molecules against various targets was distinguished by calculating the similarity between query molecules and reference molecules. For fair testing, we adopted the approach used in previous studies, selecting ligands provided in the official PDB templates as reference molecules. Notably, in scenarios assessing virtual screening capabilities, LIT-PCBA presents a more challenging task compared to DUD-E, primarily because for each target in DUD-E, the ratio of active to inactive molecules is relatively fixed with considerable structural variation. However, such “standardized” tasks as seen in DUD-E are rare in drug discovery. In contrast, there are typically challenging tasks like LIT-PCBA, especially given the drug activity cliffs, complex mechanisms of action, and significant variations in the proportions of active compounds across different tasks. Therefore, LIT-PCBA is considered the main virtual screening benchmark dataset in our study.

We compared the PhenoScreen with multiple baselines, including (I) GeminiMol, which is based on the similarity comparison of molecular conformation ensembles; PhaseShape, a method based on 3D pharmacophore and shape alignment; various molecular fingerprints (including ECFP4^24^, ECFP6^24^, FCFP4^24^, FCFP6^24^, MACCS^25^, RDK^26^, AtomPairs^27^, and Topological torsion^28^) represented by ECFP4; (II) the classic molecule docking method Glide SP^29^; and (III) PLANET^30^, a drug-target affinity (DTA) prediction-based method. The results indicate that PhenoScreen performed well in both the DUD-E and LIT-PCBA tasks (Table 2 & 3 & S2 & S3). For the 102 targets in DUD-E, PhenoScreen achieved an *AUROC* over 0.9 for 32 targets and over 0.8 for 57 targets, with an average *AUROC* of 0.803, significantly outperforming other baseline methods (Table S2).

**Table 2.** *AUROC* of the diLerent models on LIT-PCBA. Bold indicates the highest performance among all methods on this target.

| Target | PhenoScreen | GeminiMol | Phase Shape | ECFP4 | Glide SP | PLANET |
| --- | --- | --- | --- | --- | --- | --- |
| ADRB2 | 0.550 | 0.480 | **0.584** | 0.552 | 0.573 | 0.390 |
| ALDH1 | 0.551 | 0.513 | 0.494 | 0.512 | **0.788** | 0.549 |
| ESR1_ago | **0.870** | 0.800 | 0.766 | 0.790 | 0.513 | 0.668 |
| ESR1_ant | 0.595 | 0.546 | 0.570 | 0.467 | 0.450 | **0.604** |
| FEN1 | **0.644** | 0.596 | 0.441 | 0.380 | 0.408 | 0.606 |
| GBA | 0.553 | 0.569 | 0.436 | 0.441 | 0.581 | **0.689** |
| IDH1 | **0.614** | 0.500 | 0.343 | 0.393 | 0.216 | 0.503 |
| KAT2A | **0.601** | 0.607 | 0.526 | 0.440 | 0.490 | 0.498 |
| MAPK1 | **0.653** | 0.621 | 0.558 | 0.534 | 0.558 | 0.612 |
| MTORC1 | **0.467** | 0.459 | 0.427 | 0.450 | 0.490 | 0.467 |
| OPRK1 | 0.585 | 0.650 | 0.498 | **0.697** | 0.528 | 0.455 |
| PKM2 | **0.701** | 0.697 | 0.592 | 0.605 | 0.669 | 0.563 |
| PPARG | **0.762** | **0.768** | 0.747 | 0.743 | 0.670 | 0.705 |
| TP53 | 0.584 | **0.608** | 0.512 | 0.435 | **0.640** | 0.587 |
| VDR | 0.643 | 0.549 | 0.420 | 0.429 | 0.462 | 0.441 |
| Average | **0.625** | 0.597 | 0.528 | 0.525 | 0.536 | 0.556 |

**Table 3.** EF1% of the diLerent models on LIT-PCBA. Bold indicates the highest performance among all methods on this target.

| Target | PhenoScreen | GeminiMol | Phase Shape | ECFP4 | Glide SP | PLANET |
| --- | --- | --- | --- | --- | --- | --- |
| ADRB2 | 17.643 | 17.643 | **18.750** | 11.762 | 5.555 | 5.882 |
| ALDH1 | 2.509 | 2.438 | 1.618 | 2.312 | **10.336** | 1.381 |
| ESR1_ago | **10.879** | 9.791 | 8.324 | 0.000 | 7.106 | 7.687 |
| ESR1_ant | 2.364 | 2.336 | 0.980 | 0.000 | 0.409 | 3.883 |
| FEN1 | **7.583** | 3.921 | 0.815 | 0.560 | 0.602 | 5.149 |
| GBA | 3.703 | 3.124 | 3.029 | **4.907** | 4.209 | 3.011 |
| IDH1 | **5.127** | 2.563 | 0.000 | 5.127 | 0.140 | 2.564 |
| KAT2A | 1.579 | 0.000 | 2.590 | 0.523 | 1.507 | **3.108** |
| MAPK1 | **2.295** | 1.974 | 2.278 | 0.990 | 1.289 | 1.297 |
| MTORC1 | 1.030 | **5.150** | 0.000 | 1.030 | 0.842 | **2.060** |
| OPRK1 | 0.000 | 4.165 | 0.000 | 4.165 | **4.166** | 4.165 |
| PKM2 | 7.873 | 8.605 | 4.403 | **8.788** | 4.020 | 1.831 |
| PPARG | **31.114** | 26.669 | 22.798 | 22.224 | 3.879 | 3.660 |
| TP53 | **7.634** | 6.107 | 5.063 | 1.527 | 2.369 | 2.500 |
| VDR | **5.158** | 3.986 | 2.152 | 2.817 | 0.433 | 1.021 |
| Average | **7.099** | 6.565 | 4.853 | 4.449 | 3.124 | 3.280 |

Fig. 4 illustrates the visualization of molecular encoding by PhenoScreen compared to ECFP4 across various targets, showing that PhenoScreen significantly distinguishes between active and inactive molecules in feature space, with most active molecules closely matching the reference molecules in distribution, whereas some active molecules encoded by ECFP4 are further away, indicating that PhenoScreen effectively captures the distinctions in features between active and inactive molecules. For the more challenging LIT-PCBA task (Table 2 & 3 & S3), PhenoScreen’s overall screening performance and early enrichment rates surpassed those of various baseline methods in eight out of fifteen targets. Unlike Glide SP and PLANET, PhenoScreen does not utilize any target protein information, which further demonstrates its superior performance in molecular representation. Additionally, compared to GeminiMol, PhenoScreen showed higher performance in the virtual screening tasks for active molecules over most of the studied targets, suggesting that the active information in cell phenotypes indeed enhances molecular representation capability and the ability to screen active molecules.

**Fig. 4.**
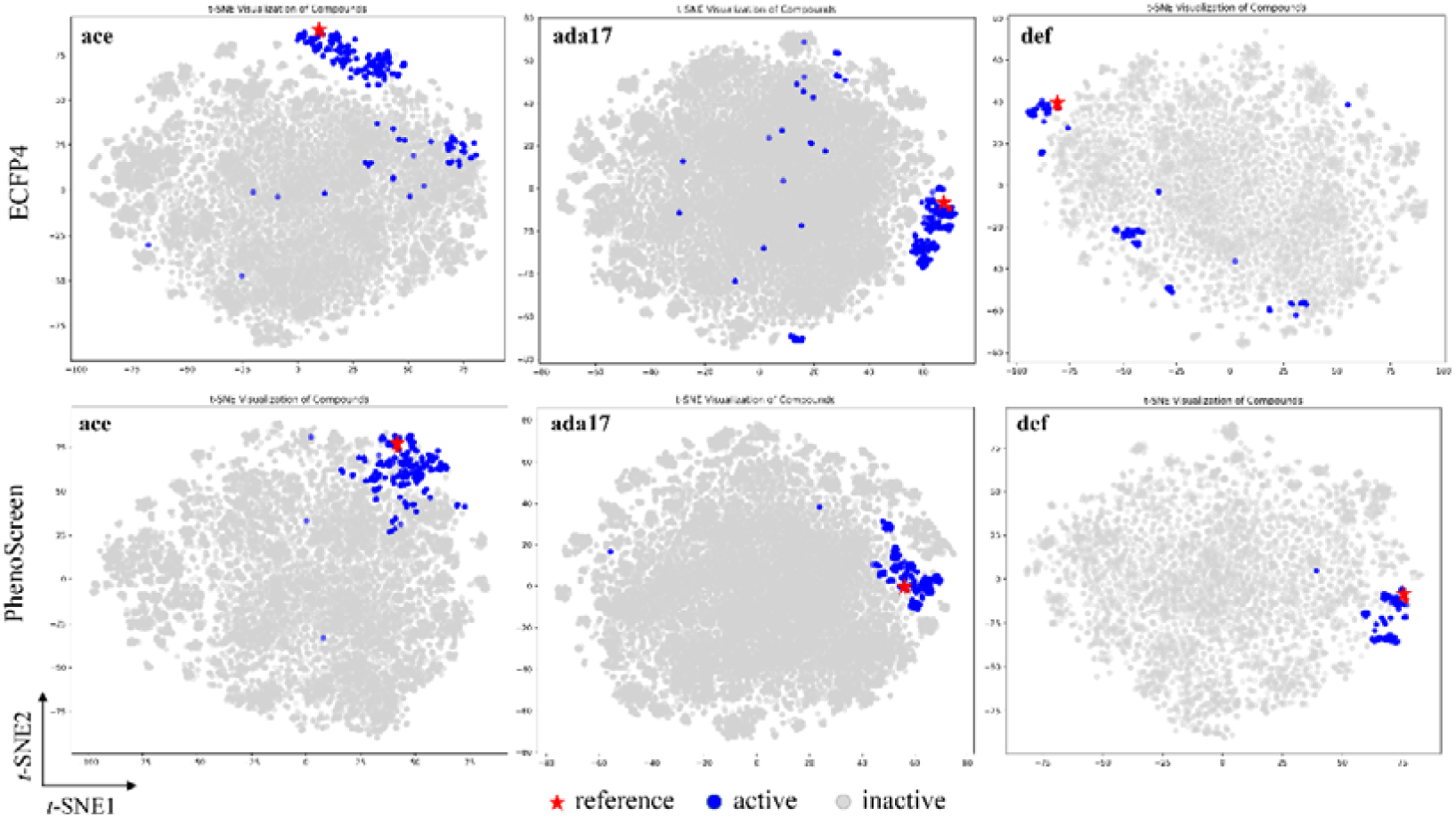
Visualization of molecular representations learned by PhenoScreen via t-SNE. The active molecules encoded by ECFP4 is further away from the reference molecule in feature space compared to PhenoScreen. This suggests that our method may be more effective in molecular representation.

### PhenoScreen Outperforms the Baseline Methods on Molecular Property Prediction

The dynamic properties of molecules, such as membrane permeability, are one of the important factors affecting drug efficacy and cellular phenotype. In the process of new drug development, the *in vivo* properties of drug molecules, including absorption, distribution, metabolism, excretion and toxicity, abbreviated as ADMET properties, are important indicators of druggability in the development of new drugs. By learning the characteristic patterns of different molecules in specific tasks, models can be established that play a crucial role in various drug design processes such as predicting molecular activity and various ADMET properties.

We compared PhenoScreen with GeminiMol,^31^ Uni-Mol,^32^ FP-GNN,^33^ various combined molecular fingerprints (CombineFP, combined by ECFP4^24^, FCFP6^24^, AtomPairs^27^, and Topological torsion^28^), and individual types of molecular fingerprints, testing 22 ADMET property prediction tasks from the TDC dataset that feature public leaderboards, including 13 classification tasks and 9 regression tasks. The results, as shown in Figs 5A and 5B, indicate that PhenoScreen’s predictive performance is comparable to Uni-Mol, which is based on large-scale pre-training using the ChEMBL database. ADMET represents a higher-level “phenotype” data, which in most cases cannot be directly linked to molecular properties from the level of cellular changes. Therefore, PhenoScreen did not show a significant improvement over the extensively pre-trained Uni-Mol in predicting ADMET properties, but it still displayed competitive performance, exceeding baseline methods in predictions of metabolism and excretion properties, and overall predictive performance on par with CombineFP and Uni-Mol. However, PhenoScreen doesn’t pre-trained at the level of hundreds of million molecules like Uni-Mol. We believe that PhenoScreen has been able to represent molecules relatively comprehensively and may show greater advantage in tasks that are more closely related to cellular phenotypes.

**Fig. 5.**
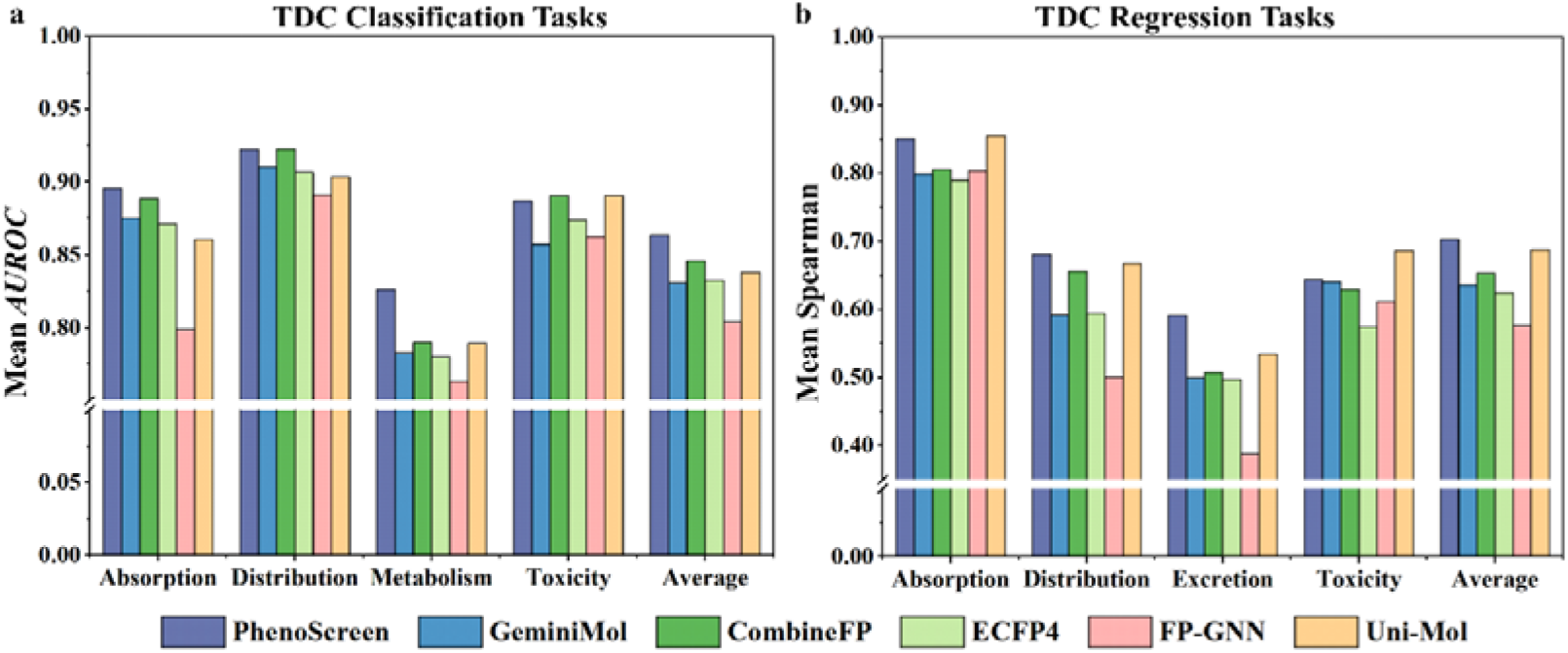
The comparison between PhenoScreen and other baseline methods in molecular properties prediction tasks. **(a)** Thirteen TDC classification. **(b)** Nine TDC regression tasks.

### Identification of Novel U2OS Inhibitors Aligned with Varying Drug-Induced Cellular Phenotypes

To assess the ability of PhenoScreen for practical drug design applications, we collected seven compounds from the Ardigen PhenAID database (https://phenaid.ardigen.com/) that can affect the phenotype of the U2OS cell line and identified four molecules from literature reports that showed inhibitory effects on osteosarcoma. With the priori information obtained from known phenotypes and activities reported in the literature, we expected to screen new molecules with similar or better phenotypes using PhenoScreen. Therefore, we performed similarity screening of these active reference molecules in the Specs compound library (https://www.specs.net/) using PhenoScreen and finally identified compounds A1-A15. Therefore, we performed similarity screening of these active reference molecules in the Specs compound library using PhenoScreen and finally identified compounds A1- A15. We measured the cellular activity of the screened compounds using the CCK8 assay (Figs 6B and 6C) and found that most of the screened molecules exhibited inhibitory activity at the cellular level. We further observed the effect of the compounds on the cellular phenotype by cell painting assay, and surprisingly, the results of the cell painting assay showed that the phenotypes of the A3, A5 and A9 were close to the reference molecules, while A11 and A15 showed higher inhibitory activity at the cellular level as compared to the reference molecules (Fig. 7). These molecules can be further investigated as potential drugs for the treatment of osteosarcoma.

**Fig. 6.**
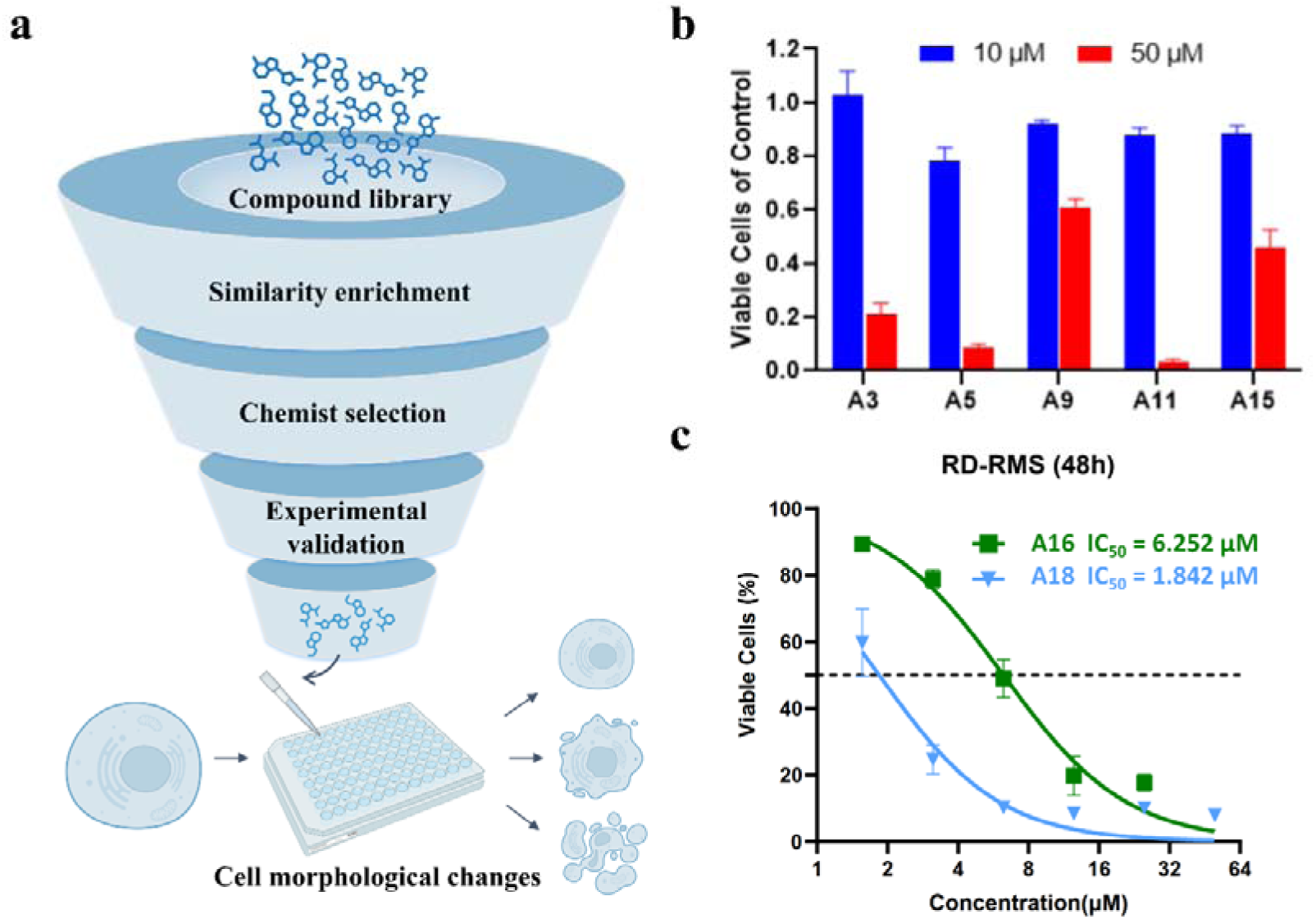
The process and experimental results of using PhenoScreen for phenotype- based drug screening. **(a)** Similarity calculations are performed between compound libraries and predefined reference molecules to enrich highly similar molecules. Chemists then select from the highly similar molecules and conduct experimental verification to observe the inhibitory ability and morphological changes of the compounds at the cellular level. **(b)** Cell viability assay results of molecules screened for U2OS. **(c)** The half-maximal inhibitory concentration (IC_50_) of the identified bioactive compounds A16 (6.252 μM, CI: 5.590 μM to 6.994 μM) and A18 (1.842 μM, CI: 1.470 μM to 2.148 μM) against rhabdomyosarcoma.

**Fig. 7.**
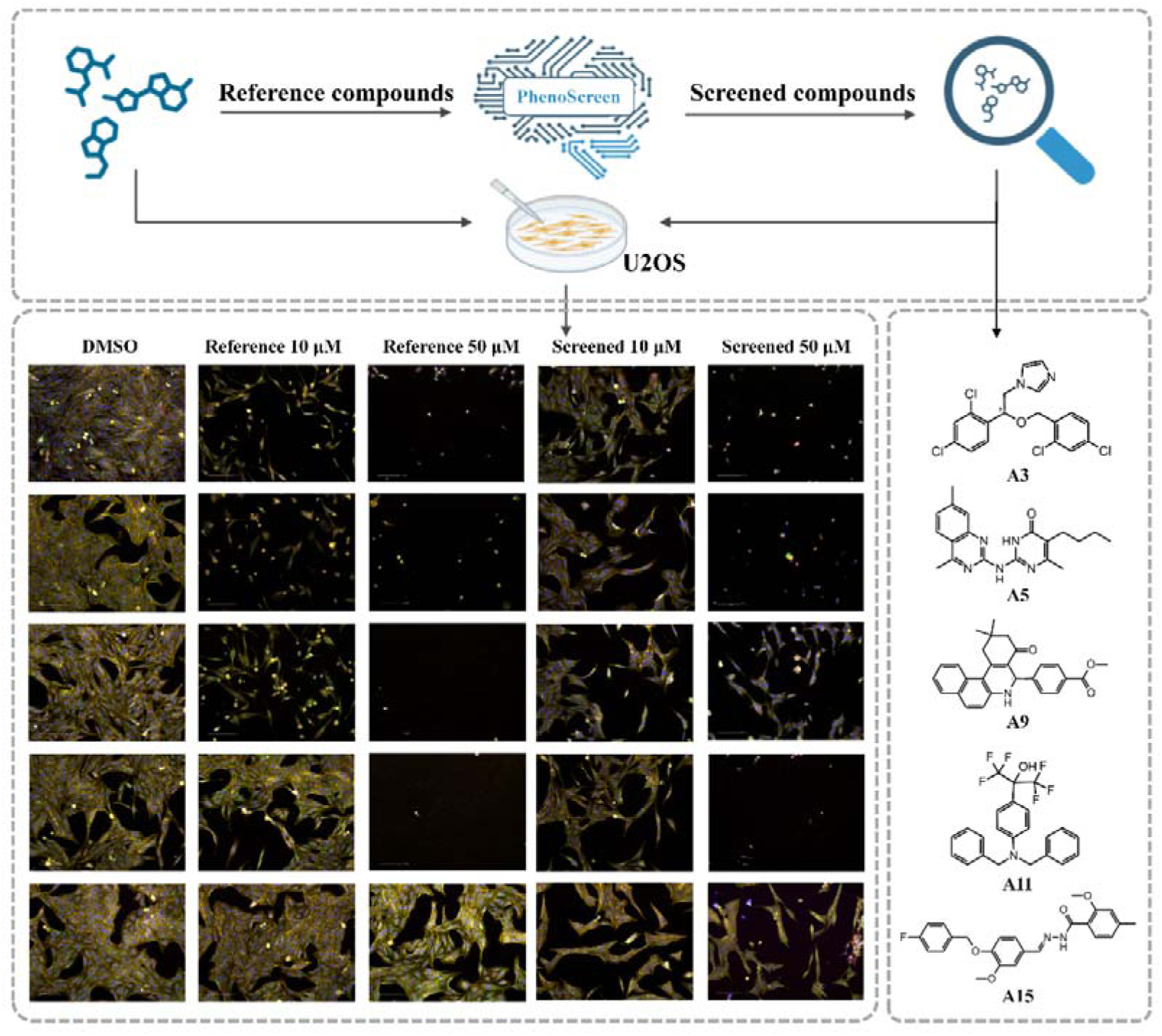
Cell phenotype results of cell painting experiments for molecules screened in the U2OS cell line showed that phenotypic changes induced by A3, A5 and A9 were similar to those of the reference molecules. In contrast, A11, and A15 exhibited more pronounced changes in cellular phenotypes compared to the reference molecules, indicating higher potential inhibitory activities.

### Exploring PhenoScreen ’ s Capability to Identify Anti-Tumor Inhibitors Across Diverse Tumor Cell Phenotypes

Furthermore, we investigated whether the screening capabilities of PhenoScreen could be generalized to diseases other than osteosarcoma. Based on preliminary studies, we identified ML324 from the literature as a potential therapeutic agent for rhabdomyosarcoma. Using PhenoScreen, we performed a similarity screen in the Specs compound library, which led to the identification of compounds A16-A20. The Cell painting results showed that these molecules have a stronger inhibitory ability compared to ML324 (Fig. 8). These results suggest that molecules such as A16 and A18 have potential as therapeutic drugs for rhabdomyosarcoma, demonstrating the generalizability of PhenoScreen for active molecule screening.

**Fig. 8.**
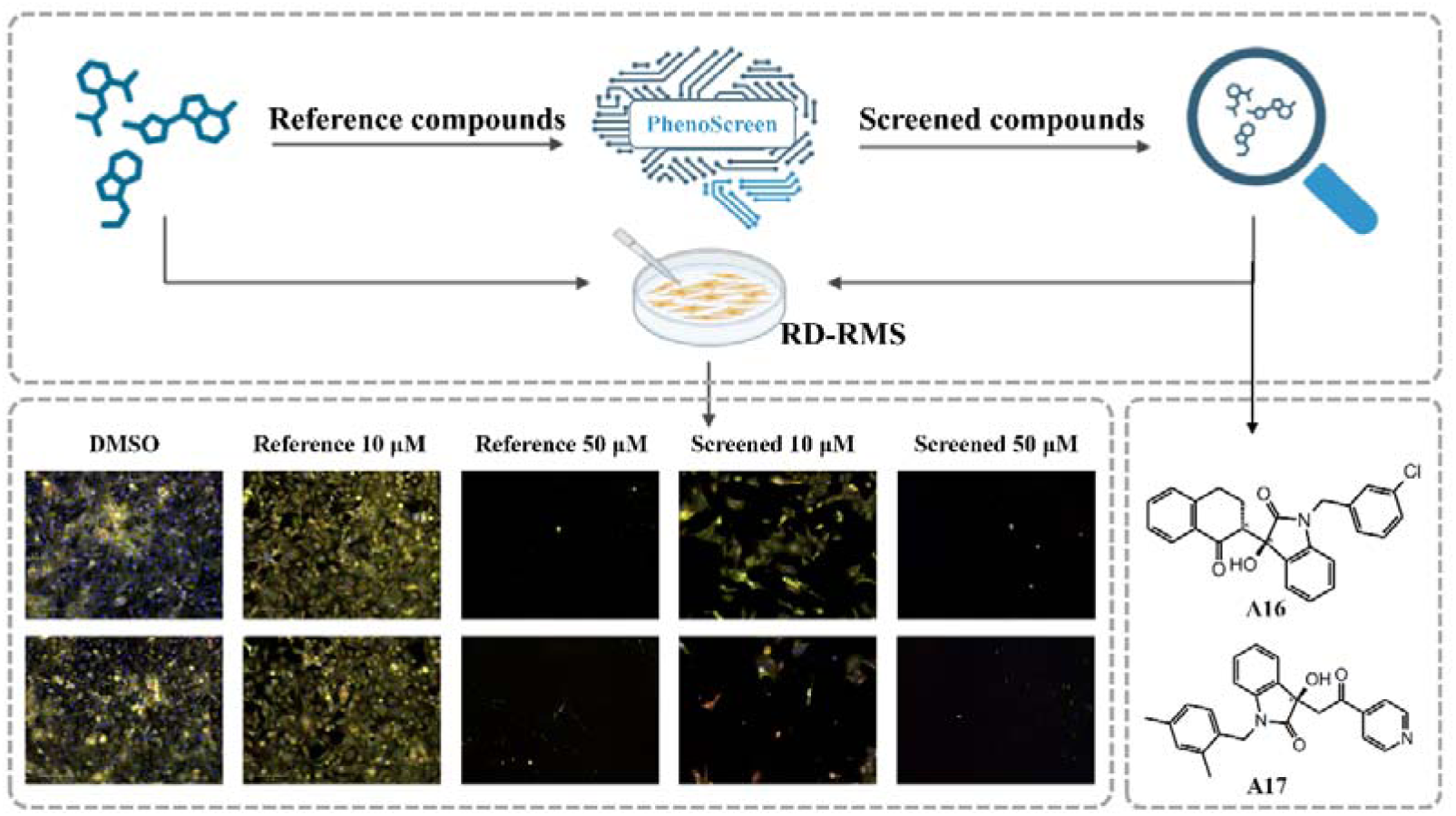
Cell phenotype results of cell painting experiments for active molecules screened for rhabdomyosarcoma. A16 and A18 showed more strong changes in cellular phenotypes compared to the reference molecules, indicating higher potential inhibitory activity.

In summary, PhenoScreen can identify potential molecules with similar phenotypes, and in some cases, even stronger inhibitory activity, by capturing the phenotypic information of molecules and their perturbations. Thus, PhenoScreen serves as an excellent hit discovery method that can be effective across various diseases. In practical applications, since reference molecules often undergo modifications to enhance activity, PhenoScreen leverages both structural and phenotypic information to further improve activity and optimize the pharmacokinetic properties of molecules. Additionally, PhenoScreen can be applied to scaffold hopping of active molecules, thereby explore more new active molecules.

### Web Server Implementation

To facilitate the implementation of the proposed PhenoScreen model by researchers, an easy-to-use web server has been established at https://bailab.siais.shanghaitech.edu.cn/services/PhenoScreen/. The web server allows users to simply upload the SMILES string of a compound of interest and select the desired molecular properties for prediction. The website then returns the molecular graph of the compound along with the predicted properties. Additionally, the server supports a compound similarity screening feature. Users can upload the SMILES string of an active compound of interest, choose from supported molecular libraries, and the website will return the top three thousand molecules ranked by similarity and their scores.

Considering the diversity of molecular structures in the molecular library and the complexity of synthesis, we provide several molecular libraries such as Specs, Chemdiv, and Enamine. Moreover, analyzing the target mechanisms of small molecules identified through phenotype-based screening can enrich the strategies for discovering new targets. Therefore, we have constructed a benchmark dataset called TIBD (Target Identification Benchmark Datasets), based on previous processing of the BindingDB database. This dataset can help researchers identify potential targets of query molecules by searching for known active compounds similar to the query molecules and determining their targets. We believe that these two applications of PhenoScreen will assist in advancing more drug discovery research.

## Conclusion

In this study, we proposed a dual-space based phenotypic drug discovery method, named PhenoScreen, which integrates the information from molecular structure space and bioactive phenotypic space. PhenoScreen not only fully characterizes the conformational information of molecules but also incorporates the active information embedded in cell phenotypes. The benchmark results presented in this paper demonstrate that PhenoScreen’s representation of molecules plays a significant role in virtual screening tasks. It performs exceptionally well in identifying active molecules, both in single-target tasks such as DUD-E and LIT-PCBA, and in various NCI cell line bioassays based on multi-target activity tests. Additionally, PhenoScreen also exhibits performance comparable to other specialized models in molecular property prediction tasks, demonstrating its potentials in extensive applications. Of course, we must acknowledge that there is substantial room for improvement in PhenoScreen. On one hand, compared to the vast chemical space, PhenoScreen has been trained using only 30,000 molecule-image pairs, which may not be sufficient for tasks requiring a comprehensive consideration of various molecular structures, such as molecular property prediction. Inevitably, it may perform slightly worse in related benchmark tests compared to extensively pre-trained molecular representation models like Uni- Mol. On the other hand, the information contained in cell phenotype data may not fully support PhenoScreen in tasks that require the representation of higher- dimensional molecular information. In future explorations, it might be possible to enhance PhenoScreen’s representational capabilities by incorporating data such as the dynamics of molecules in organoid systems. Additionally, cell image data inevitably contains noise due to experimental errors, and how to obtain cleaner and richer phenotypic data is one of the urgent problems to be solved in all related research.

It is noteworthy that changes in transcriptomic information are also crucial in phenotype-based drug design. In future research, interacting molecular, cell image, and transcriptomic information may further enhance the model’s ability to represent molecules. Additionally, care must be taken not to compromise the original encoding ability of the molecular Encoder when integrating phenotypic information into the molecular representation model, hence strategies like joint training may be required to mitigate potential risks, especially for such multimodal models. We plan to further enhance PhenoScreen’s representational capabilities in our subsequent studies. We believe that with the addition of more task-related training data, PhenoScreen could exhibit superior performance in common drug design tasks such as virtual screening and property prediction, and play a significant role in areas like genetic-chemical perturbation association modeling and phenotype-based molecule generation. This would provide researchers with valuable tools to accelerate the drug discovery process.

## Materials and methods

### Collection of Datasets

In this study, we used the cpg0012 dataset released by the Broad Institute for molecule-image contrastive learning.^21^ This dataset comprises paired perturbation molecules and microscope images from experiments conducted on 406 384-well plates. For each well, six different viewpoints of cell microscope images were taken, resulting in a total of 30,616 different perturbation molecules and 919,265 five- channel cell microscope images captured from human osteosarcoma cells (U2OS). After conducting data quality control and excluding control group images, the dataset refined to 30,611 molecules and their corresponding 757,934 microscope images with a resolution of 692 × 520 pixels. To streamline subsequent processing, all cell images were stored in npz format and divided into training, validation, and test sets with an 8:1:1 split, ensuring that multiple images from the same sample were kept within the same set.

For molecule-molecule contrastive learning, we employed a dataset from our previous study, which includes 8,000,000 pairs of intermolecular conformational space similarity (CSS) metrics used for training the molecule graph encoder.^31^

### Target-based Virtual Screening Benchmark

DUD-E and LIT-PCBA are two commonly used compound screening datasets in drug design, utilized to evaluate and test the performance of virtual screening algorithms^22,23^. DUD-E is a dataset extensively used for benchmarking molecular docking and virtual screening algorithms, covering a variety of different biological targets^22^. It includes a large number of active compounds and corresponding inactive compounds (decoy molecules). These decoy molecules are specifically designed to have similar physical properties to the active compounds, but are chemically distinct enough to ensure that the testing algorithms can differentiate between active and inactive compounds. LIT-PCBA is a large compound library containing activity test results from the PubChem BioAssay database^23^. This dataset includes multiple different bioactivity tests, providing researchers with extensive resources for cheminformatics and virtual screening studies. Unlike the intentionally designed negative compounds in DUD-E, the inactive molecules in LIT-PCBA come from experimental test results, presenting greater classification challenges. Therefore, testing the capability of screening algorithms in LIT-PCBA is a more challenging task. Due to their scale and diversity, both datasets are frequently used to test and improve the effectiveness of virtual screening technologies in drug discovery.

To ensure fair testing, we follow the approach adopted in previous studies, using ligands provided in the official PDB templates as reference molecules.^30,31^ For each target in the DUD-E dataset, there is only one reference molecule, with a positive to negative sample ratio of about 1:50; for LIT-PCBA, there may be multiple reference molecules, with a positive to negative sample ratio of about 1:1000. In the benchmark testing for LIT-PCBA, each target has several lead compounds that can serve as reference molecules. We calculate the *Pearson* similarity between all reference molecules and the query molecules, and select the highest similarity value as the final score for each query molecule.

For virtual screening tasks, we select *AUROC* (Area Under the Receiver Operating Characteristic curve) and EF1% (Enrichment Factor at 1%, Eq. (1)) as evaluation metrics.

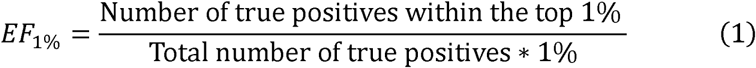

*AUROC* is an important index for evaluating the generalization performance of models, focusing on overall screening performance, while EF1% specifically targets the early enrichment rate, which is crucial for identifying highly effective compounds early in the screening process.

### Phenotype-based Virtual Screening Benchmark

We have collected 73 cell phenotype datasets from the National Cancer Institute Development Therapeutics Program (NCI/DTP) as a multi-target virtual screening benchmark test set. These datasets contain molecular activity test results from 10 different cell lines, including breast cell line (8 datasets: NCI_BT-549, NCI_HS578T, NCI_MCF7, NCI_MDA-MB-231/ATCC, NCI_MDA-MB-435, NCI_MDA-N, NCI_T-47D and NCI-ADR-RES), central nervous system cell line (8 datasets: NCI_SF-268, NCI_SF-295, NCI_SF-539, NCI_SNB-19, NCI_SNB-75, NCI_SNB-78, NCI_U251 and NCI_XF498), colon cell line (9 datasets: NCI_COLO205, NCI_DLD- 1, NCI_HCC-2998, NCI_HCT-116, NCI_HCT-15, NCI_HT29, NCI_KM12, NCI_KM20L2 and NCI_SW-620), leukemia cell line (8 datasets: NCI_CCRF-CEM, NCI_HL-60(TB), NCI_K-562, NCI_MOLT-4, NCI_P388, NCI_P388/ADR, NCI_RPMI-8226 and NCI_SR), melanoma cell line (9 datasets: NCI_LOX-IMVI, NCI_M14, NCI_M19-MEL, NCI_MALME-3M, NCI_SK-MEL-2, NCI_SK-MEL-28, NCI_SK-MEL-5, NCI_UACC-257 and NCI_UACC-62), non-small cell lung cell line (11 datasets: NCI_A549/ATCC, NCI_EKVX, NCI_HOP-18, NCI_HOP-62, NCI_HOP-92, NCI_LXFL529, NCI-H226, NCI-H23, NCI-H322M, NCI-H460 and NCI-H522), ovarian cell line (6 datasets: NCI_IGROV1, NCI_OVCAR-3, NCI_OVCAR-4, NCI_OVCAR-5, NCI_OVCAR-8, NCI_SK-OV-3), prostate cell line (2 datasets: NCI_DU-145 and NCI_PC-3), renal cell line (10 datasets: NCI_786-0, NCI_A498, NCI_ACHN, NCI_CAKI-1, NCI_RXF393, NCI_RXF-631, NCI_SN12C, NCI_SN12K1, NCI_TK-10 and NCI_UO-31), and small cell lung cell line (2 datasets: NCI_DMS114 and NCI_DMS273). The average number of test molecules per dataset is 43k, and the average number of active molecules is 2.8k. Given that the results of bioassays may be influenced by synergistic effects across different targets, and multiple active molecule data are often considered in the actual drug development process, we randomly select 50 molecules as reference active molecules for each bioassay. We use the same benchmark testing approach as adopted with LIT-PCBA.

### Molecular Property Predictions Benchmark

Establishing predictive models for molecular properties through molecular representation learning is a crucial component of ligand-based drug design methods. To further evaluate the representation capabilities of PhenoScreen, we conducted experimental tests on multiple molecular benchmarks from the Therapeutics Data Commons (TDC).^34^ TDC is a large-scale collection of chemical datasets specifically designed to advance the development of machine learning methods in drug chemistry, materials science, and biology. These datasets cover various types of chemical data, such as the ADMET (Absorption, Distribution, Metabolism, Excretion, and Toxicity) molecular properties datasets. To ensure the generalizability of molecular representation methods, the training, validation, and testing sets for all molecular property modeling tasks are divided based on molecular scaffolds. For all datasets from TDC, we use the default scaffold splitting method provided by TDC, reserving 20% of the data as the test set. Since ADMET properties, compared to molecular activity, represent a higher scale of “phenotype” that often requires consideration of more molecular structural information, this task poses a greater challenge for molecular representation methods that have not utilized extensive molecular pre-training.

For molecular property prediction tasks, *AUROC* and the Spearman correlation coefficient are chosen as the evaluation metrics for classification and regression tasks, respectively. *AUROC* assesses the ability of the model to correctly rank compounds based on their likelihood of having a desired property, which is vital for classification. The Spearman correlation coefficient measures the strength and direction of a monotonic relationship between predicted and actual values in regression tasks, providing a non-parametric measure of rank correlation.

### The Design of Dual-space Contrastive Learning Strategy

The core of PhenoScreen is a carefully designed dual-space contrastive learning representation model, which integrates molecules chemical space with their perturbated cellular morphology space, as illustrated in Fig. 1A. This representation model consists of four main modules: two feature extraction modules, the Molecule Graph Encoder and the Cell ViT Encoder, which capture information from molecular structures and cell painting images, respectively; a primary molecule-molecule contrastive learning module to establish relationships within the chemical space of different molecules; and a step-wise contrastive learning module that links a molecule’s chemical space with its associated cellular morphology space. This representation model can be then seamlessly integrated with a variety of drug design tasks, including zero-shot tasks such as virtual screening for active compounds, ADMET prediction, and other related applications.

### Molecular Graph Encoder

The Molecule Graph Encoder employs a four-layer Weisfeiler-Lehman Network (WLN) to extract molecular features, building on our previous work with GeminiMol,^31,35^ a hybrid contrastive learning framework that incorporates conformational space similarities into molecular representation learning for ligand-based drug design tasks. The Weisfeiler-Lehman (WL) algorithm enhances graph representations by iteratively refining the molecular graph through the aggregation of information from neighboring nodes (atoms) and edges (bonds). This process distinguishes between non-isomorphic graphs, which is crucial for capturing both local and global structural information and for accurately representing molecules with similar topologies but different chemical properties.

The process begins by converting SMILES (Simplified Molecular Input Line Entry System) notation into a graphical representation, which is then processed by the WLN. The molecular encoder consists of four WLN layers, with the node dimension set to 2048. After processing by the WLN, the 2,048-dimensional features undergo further processing by a readout function. This function is designed to aggregate the node features into a single vector that represents the entire molecule. This vector captures the essential molecular features necessary for downstream tasks such as virtual screening and molecular property prediction.

The use of WLN allows for effective learning of the molecular graph’s topology and atom-level interactions, making it a robust choice for capturing complex molecular characteristics essential in drug discovery and other chemical informatics applications.

### Cell ViT Encoder

The Cell ViT Encoder, based on the Vision Transformer (ViT), is designed to extract information from cell painting images.^36^ ViT excels in capturing cell images by utilizing self-attention mechanisms to process entire images, enabling it to grasp both global context and intricate cellular morphology with high flexibility. Its adaptability, scalability, and ability to leverage pretraining from large datasets make it highly effective for biological applications, such as phenotype classification and multimodal learning in drug discovery. The encoder’s parameters were initially pretrained on cellular type classification and further optimized using quadrilateral attention.

We randomly cropped each 520*696 pixels cell image to 512*512 pixels and applied data augmentation techniques such as rotation before inputting them into QFormer for feature extraction. QFormer aims to address the limitations of traditional window-based attention mechanisms in handling objects of varying sizes, shapes, and orientations. Specifically, QFormer employs a novel Quadrangle Attention (QA) mechanism, which uses an end-to-end learnable quadrangle regression module to predict transformation matrices. These matrices convert the default window into a target quadrangle for token sampling and attention computation. By leveraging QFormer, we aim to mitigate the impact of dispersed effective information and noise in cell images on feature extraction.

We first utilized a classification task to train the ViT model based on QFormer to learn the correspondence between different cell images and various drug molecules. This classification task was designed using cell images processed with 30K different drug molecules collected in the cpg0012 dataset. We designed the network parameters based on QFormer-tiny and set the patch size to 8 to avoid padding zeros in the 512*512 images. Finally, we achieved an *ACC*-1 of 26.1% and an *ACC*-5 of 53.0% on the 30K -class task. After training a robust feature extractor, we froze the main parameters of the ViT model, using it as the image encoder in subsequent contrastive learning tasks.

### Molecule-Image Contrastive learning

Given the fundamental differences between chemical and cellular spaces, and the large number of parameters in the respective encoders, the molecular encoder requires substantial weight updates to align its embeddings with those of the cellular images. This can lead to network instability and degraded learning performance. To maintain stability, we fixed most of the weights in the Cell ViT image encoder, limiting variations in image embeddings. During the contrastive learning stage between molecules and cell images, each molecule-image pair was transformed into a 2,048- dimensional embedding, with InfoNCE loss applied to minimize the distance between the embeddings of positive samples in the feature spaces.

We employed a contrastive learning strategy to align features extracted from molecular structures and cell images. Contrastive learning enhances the model’s ability to differentiate by optimizing the similarity between positive sample pairs and maximizing the differences between negative sample pairs. In this study, we utilized the InfoNCE loss function (Eq. (2))),

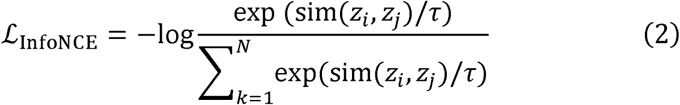

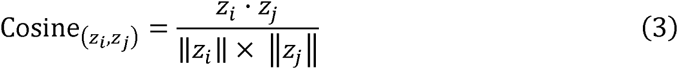

where sim(zi, zj) denotes the similarity measure between features zi and zj, typically calculated using cosine similarity (Eq. (3)). The temperature parameter ris used to adjust the smoothness of the probability distribution, and N represents the number of negative samples.

In each batch, the image features and molecular features initially correspond one- to-one. During the training process, these features are cross-paired to generate positive and negative samples. To create training samples, this study randomly selects half of the categories as positive samples, while the remaining categories are considered negative samples. Additionally, we implement a gating mechanism where the 2048-dimensional cell image features, encoded by a ViT, serve as feature weights. After normalization with a sigmoid function, these weights are multiplied with the molecular feature vectors to produce gated molecular feature vectors. These gated features are then used in contrastive training with the cell image features.

### Molecule-Molecule Contrastive learning module

To improve the efficiency of integrating chemical and cellular phenotypic spaces, we introduced molecule-to-molecule contrastive learning. This approach captures molecular interactions by leveraging similarities in chemical space and potential bioactivity, adding a constraint that allows the molecular encoder to update its weights while preserving its ability to learn conformational space similarities. For molecule- to-molecule contrastive learning, we inherited the framework from GeminiMol, which incorporates 3D conformational ensembles and pharmacophore characteristics into the embeddings, effectively linking molecular structure with bioactive information. Ultimately, this strategy achieves balanced and stable weight updates between the two networks.

The molecular graph encoder consisted of 4 WLN graph layers, and the node dimensionality was set to 2048. Molecules are converted into molecular graphs and then subsequently encoded independently. The projection head of the module employed in our study was designed to transform the 2048-dimensional encoding into meaningful CSS predictions. The network consists of a series of fully connected layers with non-linear activation functions and additional normalization to improve its learning ability. Two encoding vectors of query and reference molecules were inputted to the projection heads of CSS descriptors.

### Cell culture

The U2OS human osteosarcoma cell line (RRID: CVCL_0042) was cultured in DMEM (Meilunbio, #MA0212) supplemented with 10% fetal bovine Serum (ExCell Bio, #), 1% penicillin/streptomycin (Gibco, #15140122), and 0.1% mycoplasma prevention reagent (TransGen, #FM501). The cells were maintained at 37 °C under 5% CO_2_ in a humidified atmosphere. The RD human Rhabdomyosarcoma cell line (RRID: CVCL_1649) was cultured in DMEM (Meilunbio, #MA0212) supplemented with 10% fetal bovine Serum (ExCell Bio, #), 1% penicillin/streptomycin (Gibco, #15140122). The cells were maintained at 37 °C under 5% CO_2_ in a humidified atmosphere.

### Cell viability assay

The U2OS and RD human Rhabdomyosarcoma cells were seed at a concentration of 5000 cells/100 μL and 4500 cells/100 μL per well into 96-well plates, respectively. After 24 h incubation, the cells were exposed to DMSO or the compounds with a final DMSO concentration of 0.2%, and cultured for another 48 h. Cell viability was assessed using Cell Counting Kit-8 (CCK8) (TargetMol, #C0005). The absorbance was determined at 450 nm with the EnSpire Multimode PlateReader (PerkinElmer). The percentage of surviving cells was calculated from the absorbance ratio of compounds treated to untreated cells.

### Cell Painting and high-content imaging

The Cell Painting staining was performed according to a modified protocol by Bray et al. using PhenoVue Cell Painting kit (Revvity, #PING11). Briefly, cells were seeded in 384-well microplates (Coring, #3764BC) at a density of 2000 cells/30 μL per well. The culture medium was replaced with 40 µL of 2% FBS in DMEM with DMSO or compounds 24 h after seeding. The final DMSO concentration was 0.2%. After a 48h treatment, the PhenoVue 641 Mitochondrial Stain (500 nM final concentration) was added. The plates were incubated for 30 min in the dark at 37 °C. Cells were then fixed using 16% paraformaldehyde (Alfa Aesar, #043368.9L; 4% final concentration) at RT for 20 min, followed by four washes with 1× HBSS (Gibco, # 14065056). Subsequently, cells were permeabilized with 0.1% Trition X-100 and stained with a staining solution containing the following dyes at their respective final concentrations: PhenoVue Fluor 555-WGA (43.7 nM), PhenoVue Fluor 488-Concanavalin A (48 nM), PhenoVue Fluor 568-Phalloidin (8.25 nM), PhenoVue Hoechst 33342 Nuclear stain (1.62 µM), PhenoVue 512 Nucleic acid stain (6 µM) in the 1× PhenoVue Dye Diluent A (HBSS + 1% BSA). After a final incubation period of 30 min at RT, the plates were washed four times with 1× HBSS prior to imaging.

Cell imaging was conducted using the Operetta High Content Imaging System (PerkinElmer) equipped with a 20× water-immersion objective. Five channels were employed to visualize the various fluorescent stains: DNA (excitation: 360-400 nm; emission: 410-480 nm), ER (excitation: 460-490 nm; emission: 500-550 nm), RNA (excitation: 460-490 nm; emission: 560-630 nm), AGP (excitation: 560-580 nm; emission: 585-605 nm), and Mito (excitation: 620-640 nm; emission: 650-760 nm). Four fields of view were captured per well.

## Supplementary Information

**Table S1** Performance of each method on each subtask in the NCI dataset.

**Table S2** Performance of each method on each subtask in the DUD-E.

**Table S3** Performance of each method on each subtask in the LIT-PCBA.

## Availability of data and materials

We have released the datasets model weight used in this study which are publicly available at Zenodo (https://zenodo.org/records/13943032) and Hugging Face (https://huggingface.co/Sean-Wong/PhenoScreen). The cell painting dataset for model training is available at: http://gigadb.org/dataset/100351.^21^ All the source code are publicly available through the GitHub (https://github.com/Shihang-Wang-58/PhenoScreen).

## Author contributions

S.W., L.W, S.G, and F.B. contributed to conceptualization. S.W., W.Q., L.W., and Y. Z. contributed to model development. S.W., L.W., and R. L. contributed to benchmark test. S.W., P.R., Y. Z., Z. L., and W.Z. contributed to data curation. Q.H. and J.Y. contributed to experiment test. Y.T. developed the web server. S.W., S.G, and F. B. contributed to original draft writing. All authors contributed to review and edit the manuscript.

## Acknowledgements

This work is supported by ShanghaiTech AI4S Initiative SHTAI4S202404. The authors appreciate the technical support provided by the engineers of the high- performance computing cluster of ShanghaiTech University. The authors thank the Discovery Technology Platform at Shanghai Institute for Advanced Immunochemical Studies, ShanghaiTech University for instruments and technical support regarding High-content imaging experiments.

